# Home range size and population density are negatively correlated in wild felids globally

**DOI:** 10.64898/2026.05.16.725626

**Authors:** Noé Bugaud, Stefano Anile, Arthemis Moraru, Sébastien Devillard

## Abstract

**Aim:** Home range size is a fundamental aspect of animal spatial ecology, and understanding the factors that shape it is important for conservation purposes. Several hypotheses, based on energy needs or competition, assume that home range size negatively correlates with population density. However, this pattern has been little investigated on a global scale, and it remains unclear whether it would stand at both intra- and interspecific levels. To fill this gap, we conducted a global exploration of this relationship at the level of an animal family.

**Location:** Global.

**Time period:** Contemporary.

**Major taxa studied:** Wild *Felidae*.

**Methods:** Individual home range size records (*n* = 1022) and population density estimates (*n* = 1061) were retrieved from the literature for 23 felid species across the world. We first investigated the interspecific relationship by modelling the median home range size of a species as a function of its median population density. To study the intraspecific relationship, we spatially merged data points based on their spatial or temporal proximity. We then applied a mixed-effects linear model using species as a random factor.

**Results:** We found that home range size was negatively associated with population density, at both interspecific (−1.323 ± 0.180, *p* < 0.001) and intraspecific levels (−0.569 ± 0.201 to - 0.537 ± 0.201 depending on the merging approach, *p* < 0.01). Landscape features were also predictors of home range size, without confounding the effect of population density.

**Main conclusions:** Several processes likely govern the relationship between home range size and population density: differences in body mass between species may drive the interspecific relationship, whereas the intraspecific pattern is probably explained by conspecific competition. Although more research is needed to quantify their relative contribution, our study highlights a worldwide ecological pattern that exists at multiple biological levels in the wild.

## Introduction

One of the main goals of animal ecology is to understand the processes that influence the life and behaviour of wild species, as well as their ability to cope with natural and anthropogenic stressors (Speakman et al., 2025). In this context, understanding animals’ spatial ecology—how they use and move through space—is essential for conserving biodiversity because it provides fundamental information like species-specific habitat requirements, migration routes, and territory sizes, which are critical for designing effective protected areas, wildlife corridors and conservation actions.

Animals roam throughout the environment to feed, mate, find shelter or care for young. The area in which they carry out these activities is referred to as the home range (Burt, 1943). Home range size (hereafter HRS) is the result of a complex energetic trade-off, which takes into account food acquisition, travel cost, competition with conspecifics, as well as predation risk and prey availability (Sells & Mitchell, 2020). It is therefore unsurprising that multiple factors can influence HRS, like sex, seasonality (Kjellander et al., 2004; Kubala et al., 2024), food abundance (Kittle et al., 2015), foraging mode (McNab, 1963), or anthropogenic activities (Oh, Moteki, Nakanish & Izawa, 2010; Šálek, Drahníková & Tkadlec, 2015).

Another crucial ecological parameter which can modulate HRS is population density (hereafter PD), since it can determine the intensity of intraspecific competition and the distribution of limited natural resources (Balluffi-Fry et al., 2025; Schoepf, Schmohl, König, Pillay & Schradin, 2015). Gaining a better understanding of the relationship between PD and HRS would prove particularly important for understanding disease transmission across different levels of PD, because transmission depends not only on PD, but also on how animals use space (Pope et al., 2007).

The relationship between HRS and PD is expected to be negative, for at least three reasons. First, energy expenditure increases with body mass (Kleiber, 1947), which implies that, in a limited environment, larger animals are assumed to have lower PDs—a pattern known as Density-Mass allometry (Damuth, 1981)— and larger HRSs to meet their energetic needs (McNab, 1963). Therefore, according to this “body mass hypothesis” (H_1_), we expect HRS and PD to negatively correlate, because they would be driven by body mass in opposite directions. Moreover, since both HRS and PD scale allometrically with body mass, the HRS-PD relationship is also expected to follow a power law and should therefore appear linear on a log-scale.

A second hypothesis (H_2_) states that PD and HRS are both influenced by a third variable, namely, resource density or, more broadly, habitat quality (Balluffi-Fry et al., 2025). Indeed, areas with abundant resources are expected to sustain higher PDs (Santini et al., 2018), while also reducing the need to roam across larger areas in search of food. Several studies have found support for this hypothesis, by showing a local negative correlation between HRS and prey density (Kubala et al., 2024; Kittle et al., 2015), or a reduction of HRS when supplemental food was provided (Schoepf et al., 2015).

A third hypothesis (H_3_) states that increasing PD induces a decrease in HRS due to intraspecific competition, notably competition for food. Some studies indeed reported an increase in HRS after removing individuals from the population (Henderson, Warren, Cromwell & Hamilton, 2000; Schoepf et al., 2015), and PD can be a better predictor of HRS than overall resource density (Balluffi-Fry et al., 2025). For non-territorial species, a postulated mechanism is that, by restricting their home range, animals concentrate their movements on their most familiar area, and thus avoid an inefficient foraging strategy when competition increases (Balluffi-Fry et al., 2025). In addition, for territorial species, the cost of defending a territory increases with the number of neighbours, thus animals are expected to shrink their home range with increasing PD (Sells & Mitchell, 2020). Assuming that territories are exclusive and that the total area covered by a population remains unchanged, we can predict that HRS would be inversely related to PD and would therefore scale proportionally to PD^-1^ (Jetz, Carbone, Fulford & Brown, 2004).

In line with these predictions, a negative relationship between HRS and PD was the most common result of studies that examined this question (Šálek et al., 2015; Aronsson et al., 2016; Kauhala & Holmala, 2011). However, although rare, a few studies did not find this pattern (Kilpatrick, Spohr & Lima, 2001; McNulty, Porter, Mathews & Hill, 1997).

In general terms, most of the research addressing the relationship between HRS and PD focused on single species, and are limited to a local scale (e.g. at the population level). A comprehensive study of the HRS–PD relationship at a higher taxonomic level is lacking, and we aim to fill this knowledge gap by conducting a global multi-species study using felids as model taxa. *Felidae* is a family of charismatic mammals with 40 extant wild species (Kitchener et al., 2017). Being carnivores, they play a major ecosystem role as top predators. Moreover, they display territorial behaviours (Bradshaw, 2016), albeit to varying degrees, and are therefore suitable models to test whether a negative correlation between HRS and PD is supported, at both inter- and intraspecific levels.

By taking advantage of two comprehensive datasets, one on PD (Azizan, Anile, Nielsen, Paradis & Devillard, 2023), and the other on HRS (Moraru, Anile & Devillard, 2026), we created geographically meaningful HRS-PD pairs following an approach similar to Azizan et al. (2023), but modified on purpose. We then modelled HRS as a function of PD, at the interspecific level using the generalised least squares (GLS) method, and at the intraspecific level using a linear mixed-effects model (LMM). We expect the relationship between HRS and PD to be negative at both interspecific and intraspecific levels, under the hypotheses mentioned above. To disentangle between H_2_ and H_3_ at the intraspecific level, our models included environmental factors thought to influence HRS (Moraru et al., 2026), from which we derived the following predictions. If H_2_ is true, we predict that habitat quality would positively covary with PD and negatively covary with HRS, whereas if it is false, one of these correlations should be absent. Under H_3_, the effect of PD should remain significant when adding environmental variables, whereas if H_3_ is false, the effect of PD should be absent when these confounding factors are taken into account. We finally discuss the causes, mechanisms and consequences of our results.

## Material and Methods

### Home range size and population density databases

We based our analysis on two independent datasets compiling data from the literature on *Felidae*: one containing HRS estimates (hereafter HR dataset; Moraru et al., 2026) and another one containing PD estimates (hereafter PD dataset; Azizan et al., 2023). All records were annotated with spatial coordinates (latitude and longitude), country of origin, and temporal information (i.e. the start and end years of the survey).

The HR dataset comprised individual records of annual HRS estimates from studies using either satellite or radio-tracking. Two methods were used for estimating HRS: the Minimum Convex Polygon (MCP) with 100% isopleth, and the Kernel Density Estimation (KDE) with 95% isopleth. As these two methods may give different estimates (Kubala et al., 2024), we decided to distinguish them in the subsequent steps of the analysis. To obtain population-level estimates, we averaged individual HRS by species, sex, HRS calculation method (MCP 100% or KDE 95%) and locality (i.e., a unique set of coordinates that could include multiple individual records).

The PD dataset comprised PD estimates from studies using camera traps. We only retained PD estimates originating from Spatially Explicit Capture Recapture (SECR; Efford, 2004) models, the gold standard for estimating PD in individually recognisable felid species.

### Relationship between HRS and PD across species

To investigate the relationship between HRS and PD at the interspecific level, we considered the median HRS and the median PD for each species present in both HR and PD datasets. We then modelled HRS as a function of PD, using R software version 4.5.0 (R Core Team, 2025). In virtue of the expected allometric link between these two variables, we log_10_-transformed them to linearise the eventual relationship. To account for non-homogeneity of the residual variance (see Figure S10.1), we used the generalised least squares (GLS) method (Zuur, Ieno, Walker, Saveliev & Smith, 2009), as implemented in the *gls* function from the ‘nlme’ package (Pinheiro et al., 2017). Two variance structures were tested: *varExp* and *varConstPower*, as a function of log_10_(PD). The structure with the lowest corrected Akaike Information Criterion (AICc) value was retained for the subsequent steps.

To assess whether the species could be considered as independent observations, we computed the phylogenetic signal using Pagel *λ* (Pagel, 1999), based on a phylogeny of *Felidae* (Li, Davis, Eizirik & Murphy, 2016), and using the *phylosig* function from the ‘phytools’ package (Revell, 2012). Finally, we checked the assumptions of the GLS model (Zuur et al., 2009) using the relevant diagnostic plots (Figure S10.1).

### Relationship between HRS and PD within species

#### Pairing HRS and PD values

To study the relationship between HRS and PD at the intraspecific level, we created pairs of HRS and PD values at the population level. However, HRS and PD were not collected at the very same sites, therefore we needed to join them in a biologically relevant manner. For this purpose, we modified the method developed by Azizan et al. (2023) in their study of the relationship between PD and genetic diversity in felids.

In a first step, HRS and PD data points were clustered based on their coordinates using the *kmeans* function. Initial centroids were given by either HRS or PD coordinates. Whenever a given cluster contained only HRS or PD estimates, making it impossible to create HRS-PD pairs, we assigned these “orphan” clusters to the nearest cluster (i.e. a PD data point belonging to an “orphan” cluster was assigned to the cluster of the nearest HRS data point, and *vice-versa*). We then computed the distance between HRS or PD data points and the centre of their respective cluster using *distm* function from ‘geosphere’ package (Hijmans, 2010a), and only retained the records for which this distance was inferior to the median dispersal range of the species. This latter value was estimated from the mean HRS of the species, using the formula: 7 * HRS^0.5^ (Bowman, Jaeger & Fahrig, 2002). Therefore, this filtering process ensured that each cluster was biologically coherent by keeping in the dataset only records close to the cluster centroid.

After this first step, we obtained clusters comprising one or several HRS and PD estimates. We then attempted to match HRS and PD values, using two approaches. The first was called “simple averaging”, and consisted in pairing the average HRS of a cluster (distinguishing by sex and HRS calculation method) with the average PD of the same cluster (Figure S1.1a).

This method had the advantage of simplicity, but it did not take into account the proximity of the sampling sites within a cluster. We thus developed an alternative method, called “spatio-temporal averaging”, in which each HRS record was paired with a *weighted* average of all the PD records in the cluster (Figure S1.1b), weights being negatively related to spatial distance and temporal delay between HRS and PD records. We chose to average PD values instead of HRS values, in part because PD estimates were more numerous than HRS estimates at the population level; this method therefore avoided data duplication.

To investigate whether or not our results were sensitive to this clustering approach above, and following Azizan et al. (2023), we also conducted a “country approach”, in which records of the same species were first grouped together based on their country, and then paired using either simple averaging or spatio-temporal averaging. This method was also used to capture patterns that could occur at very large spatial scales.

#### Environmental variables

Resource densities and human activities, including agriculture and roads, have been shown to influence HRS in mammals, whether positively or negatively (Oh et al., 2010; Šálek et al., 2015; Kubala et al., 2024, Moraru et al., 2026). To account for these potential confounding factors, and help in deciphering between the potential processes explaining the relationship between HRS and PD at the intraspecific scale (H_2_ or H_3_), we annotated each locality with various environmental variables: net primary productivity (NPP), Artiodactyla richness (AR), Rodentia richness (RR), elevation, human footprint index (HFI), human population density (HPD), road density, croplands density and pastures density. As felids mostly feed on other mammals (Lamberski, 2014), the best proxy for resource density would be mammal density. However, such data was not available on a fine spatial scale; therefore, we used AR, RR, and NPP as proxies of resource availability. All values were extracted from raster maps using the R package ‘raster’ (Hijmans, 2010b). For the variables to be representative of the area where the animals are likely to move, we considered a buffer around the locality of each HRS record, whose radius was equal to the species median dispersal range (Bowman et al., 2002), and we averaged all values in this buffer. The relevant references and the spatial resolutions of the variables are given in Table S2.1. All variables were centred and scaled before modelling.

#### Statistical analysis

To investigate the intraspecific relationship between HRS and PD, we opted for a linear mixed-effects (LMM) model, and considered species as a random factor. As above, HRS and PD were log_10_-transformed. We initially considered the following fixed effects: PD, sex, HRS calculation method, NPP, elevation, AR, RR, HFI, HPD, road density, pastures density, and croplands density. We also added an interaction between sex and PD, since males and females do not display the same territorial behaviours (Kjellander et al., 2004; Kubala et al., 2024), and might respond differently to a change in PD. To assess the collinearity between the explanatory variables, we calculated the Spearman correlation coefficient *ρ* for each pair of quantitative variables, and we further calculated the variance inflation factor (VIF) of both quantitative and categorical variables using the *vif* function from the ‘car’ package (Fox, 2001). Variables were considered collinear if *ρ* > 0.6 or VIF > 3.

We fitted models using Restricted Maximum Likelihood (REML), as implemented in the ‘lme4’ package (Bates, 2016). To account for the variability in sampling effort, we assigned to each pair *i* a weight (*W_i_*) proportional to its contribution in the model:

*W_i_* = *n_HRS(i)_* × *n_PD(i)_*

where *n_HRS(i)_* and *n_PD(i)_* are the number of individual HRS records and PD records represented in the pair *i*, after scaling (to give equal importance to HRS and PD data), subtracting the minimum value and adding 1 (to have weight values greater than or equal to 1). We chose to multiply *n_HRS_* and *n_PD_* to give more weight to “balanced” pairs, i.e. pairs with a similar number of HRS and PD estimates.

To select the most parsimonious model, *sensu* best model, we started with a full umbrella model comprising all the fixed effects, minus those that were collinear with each other. We first fitted three different versions of this full model: one with species as a random intercept, one with species as a random slope, and one with species as a random intercept and a random slope. Hence we selected the most parsimonious random structure, i.e. the one with the smallest AICc value.

Then, we selected the most parsimonious fixed structure using *dredge* function from ‘MuMIn’ package (Bartoń, 2010), which generated all the possible combinations of fixed effects and ranked them according to their AICc while keeping constant the random structure. When several equivalent competitive models were supported (ΔAICc < 2), we performed conditional averaging using *model.avg* from ‘MuMIn*’* package. For each fixed effect, the *p*-value was computed using ‘lmerTest’ package (Kuznetsova, Brockhoff & Christensen, 2017). We further quantified the variance explained by fixed effects alone by calculating the marginal coefficient of determination (R²marg) of the model having the lowest AICc value, using the *r.squaredGLMM* function from ‘MuMIn’ package. We also quantified the variance explained by both fixed and random effects by calculating the conditional coefficient of determination (R²cond). Lastly, we checked whether or not the assumptions of the linear mixed models were respected (Zuur et al., 2009) using the relevant diagnostic plots (Figure S10.2).

## Results

The PD and the HR datasets originally included 1105 and 1137 records, respectively. When focusing only on the 23 species present in both datasets, we were left with 1061 PD estimates and 1022 individual HRS records, spanning 25 countries (Table S3.1, Figure S3.1). The HRS records were then grouped by location, sex, and HRS calculation method, resulting in 292 population-level HRS estimates (Table S3.1, Figure S3.1).

### Interspecific relationship between HRS and PD

Among the 23 species of felids present in both HR and PD datasets, *Lynx pardinus* had an extremely low PD (see Figure S4.1), likely because it was critically-endangered (CR) during the monitoring (IUCN, 2025); it was thus considered an outlier and removed. When fitting a simple linear model between HRS and PD at the interspecific level, we saw a clear violation of homoscedasticity (Figure S10.1), so we opted for the GLS method. The *VarExp* variance structure was more parsimonious (ΔAICc = 2.5), and was kept for the analysis. As displayed in Figure 1, HRS and PD showed a significant negative relationship (*β* = -1.323, SE = 0.180, *p* < 0.001). No significant phylogenetic signal was observed in both datasets (*λ_HRS_* = 0.124, *p* = 0.621; *λ_PD_* = 0.119, *p* = 0.510).

**Figure 1.**
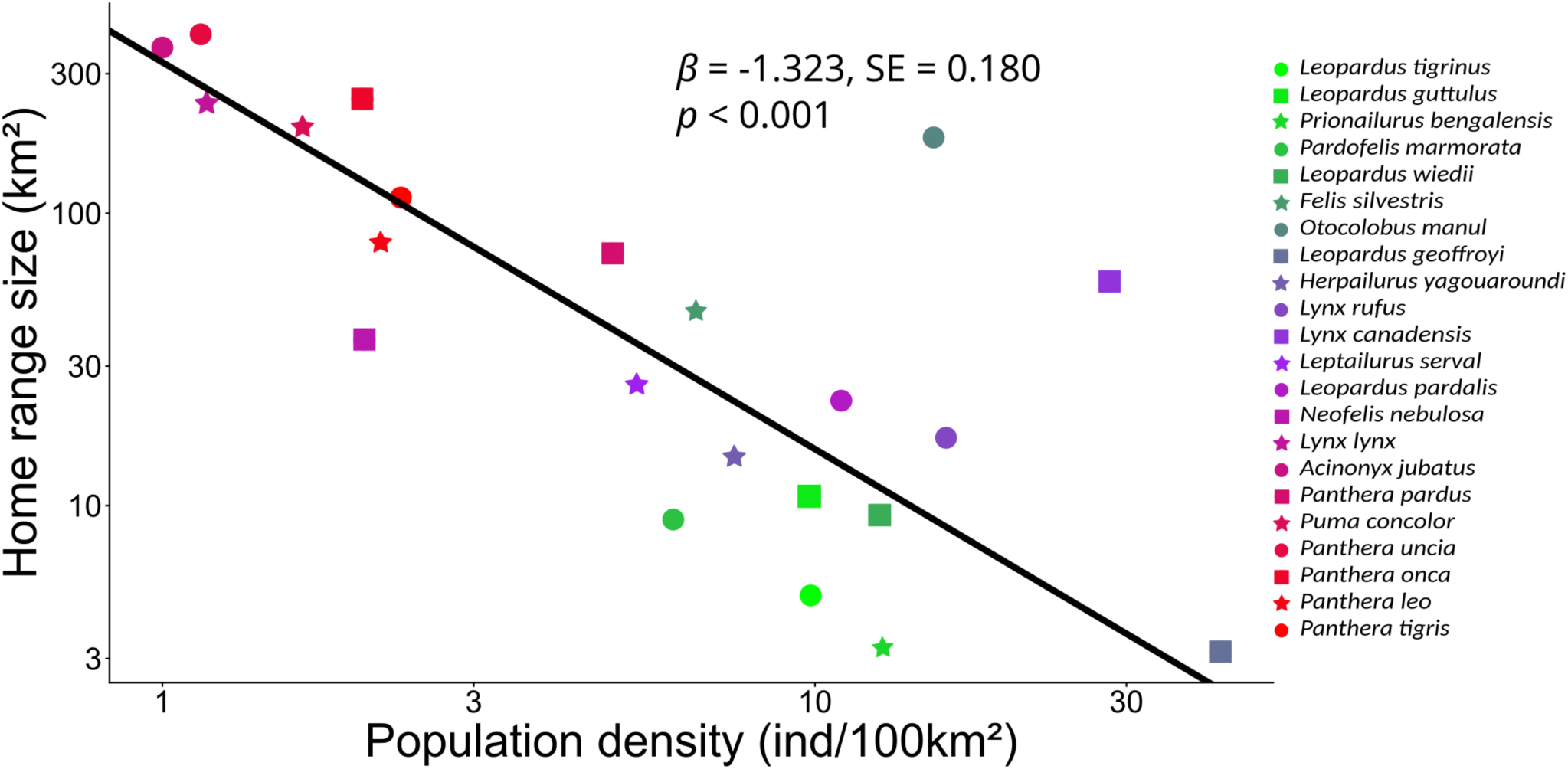
Home range size (HRS) and population density (PD) are negatively correlated across *Felidae* species. Each point corresponds to the median HRS and the median PD of a given species, considering its worldwide records. The regression line was obtained from the generalised least squares (GLS) model log_10_(HRS) ∼ log_10_(PD) using the *varExp* variance structure. SE: standard error.

### Intraspecific relationship between HRS and PD

We obtained more clusters using PD (n = 41) than HRS (n = 30) locations as initial centroids in the K-means method (Table S5.1), so we opted for the former. These 41 clusters belonged to 10 species, and enabled us to create 95 and 120 pairs following simple and spatio-temporal averaging, respectively (Table S3.1). The median time interval between HRS and PD studies (Table S7.1) was lower when using spatio-temporal averaging (5.4 years) than when using simple averaging (8.1 years).

Collinearity was found between HPD, HFI, croplands density and road density (*ρ* > 0.6, Figure S8.1). We hence kept croplands density and removed the other variables since: 1) croplands density has a more direct interpretation than HFI, which is the result of multiple human disturbance factors; 2) croplands are directly linked to the use of space, contrary to HPD; 3) croplands density generally showed higher variance across all analyses, making it more valuable for modelling purpose. Once HPD, HFI and road density were removed, the VIF of the remaining explanatory variables (including sex and HRS calculation method) was inferior to 3.

The most parsimonious random structure was the one with species as random slope and intercept (Table S5.2). We then selected the fixed structure and averaged two equivalent competitive models for each merging approach (ΔAICc < 2, Table S5.3). Those models verified the assumption of linear mixed-effects models (Figure S10.2). As shown in Table 1 and Figure 2, we found evidence of a negative association between PD and HRS, both using simple averaging (*β* = -0.537, SE = 0.201, *p* < 0.01) and spatio-temporal averaging (*β* = - 0.569, SE = 0.201, *p* < 0.01). HRS also negatively correlated with croplands density in both approaches (*β* = -0.211, SE = 0.051, *p* < 0.001 and *β* = -0.161, SE = 0.049, *p* < 0.01, respectively). Males had larger home ranges than females (*p* < 0.001), but the interaction between sex and PD was not retained. The MCP method yielded higher HRS values than KDE (*p* < 0.01). Overall, the best model explained 73% of the total variance in simple averaging (R²cond = 0.730), and 66% in spatio-temporal averaging (R²cond = 0.663). Fixed effects alone explained 42% and 37% of the total variance, respectively (R²marg = 0.420 and 0.373).

**Figure 2.**
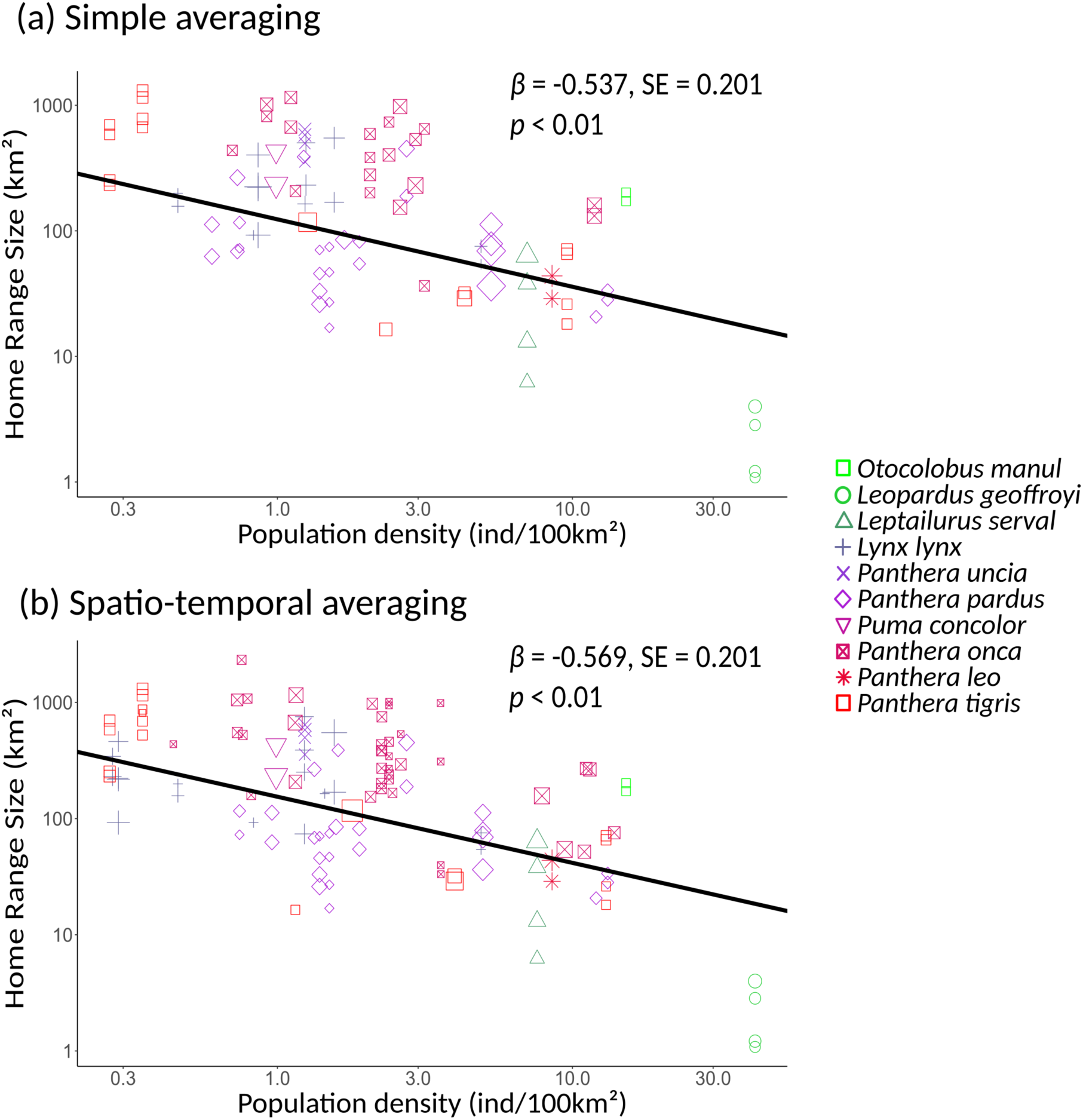
Home range size (HRS) and population density (PD) are negatively correlated within *Felidae* species. Worldwide HRS and PD estimates were paired according to their spatial coordinates using K-means clustering algorithm. **(a) Simple averaging**: in a cluster, one average PD was paired with one average HRS per sex and per HRS calculation method (n = 95 pairs). **(b) Spatio-temporal averaging**: in a cluster, HRS was paired with a weighted mean of all PD records, weights being calculated from spatial and temporal proximity between both records (n = 120). Regression lines represent the intraspecific effect of PD on HRS, obtained by linear mixed-effects modeling with species as a random intercept and a random slope. Dot size denotes the weight of a given HRS-PD pair in the model, and this weight is calculated from the number of PD and HRS records represented in the pair. SE: standard error.

**Table 1.**
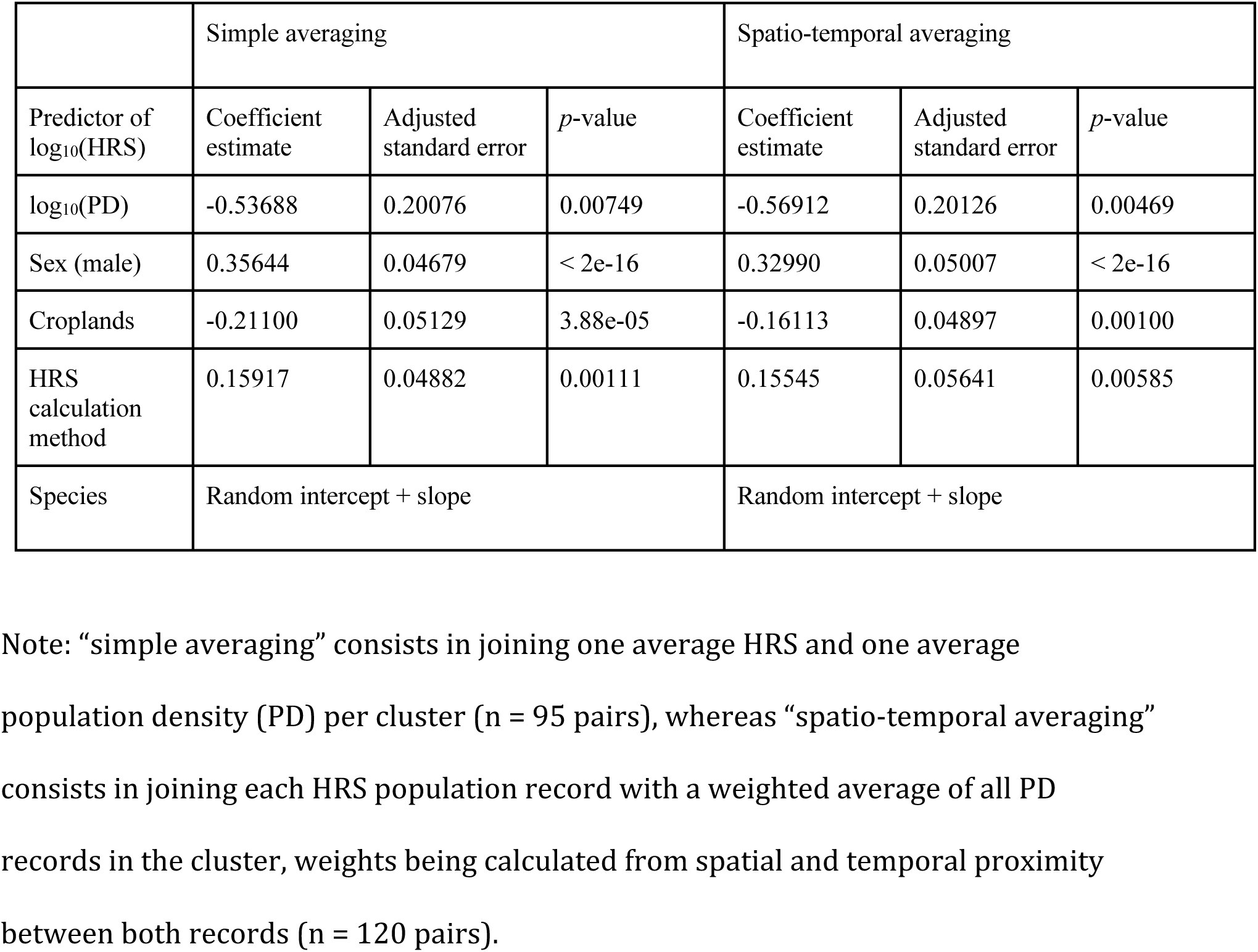
Global predictors of home range size (HRS) at the intraspecific level in *Felidae*, obtained by averaging the two best linear mixed-effects models (ΔAICc < 2).

|  | Simple averaging |  |  | Spatio-temporal averaging |  |  |
| --- | --- | --- | --- | --- | --- | --- |
| Predictor of $\log_{10}(\text{HRS})$ | Coefficient estimate | Adjusted standard error | *p*-value | Coefficient estimate | Adjusted standard error | *p*-value |
| $\log_{10}(\text{PD})$ | -0.53688 | 0.20076 | 0.00749 | -0.56912 | 0.20126 | 0.00469 |
| Sex (male) | 0.35644 | 0.04679 | $< 2e-16$ | 0.32990 | 0.05007 | $< 2e-16$ |
| Croplands | -0.21100 | 0.05129 | 3.88e-05 | -0.16113 | 0.04897 | 0.00100 |
| HRS calculation method | 0.15917 | 0.04882 | 0.00111 | 0.15545 | 0.05641 | 0.00585 |
| Species | Random intercept + slope |  |  | Random intercept + slope |  |  |

Note: “simple averaging” consists in joining one average HRS and one average population density (PD) per cluster ( $n = 95$ pairs), whereas “spatio-temporal averaging” consists in joining each HRS population record with a weighted average of all PD records in the cluster, weights being calculated from spatial and temporal proximity between both records ( $n = 120$ pairs).

## Discussion

While several single-species studies reported a negative trend between HRS and PD, a comprehensive study of this pattern at a global scale and at a higher taxonomic level was lacking. Here we first show that HRS and PD are negatively correlated between felid species. To investigate the intraspecific level, we used K-means clustering, followed by two alternative methods (simple averaging and spatio-temporal averaging) to generate HRS-PD pairs within each species. Regardless of the method, we found a consistent negative relationship between HRS and PD within species, in addition to a negative association with croplands density. Furthermore, the negative correlation between HRS and PD was also found with the country approach (Tables S6.1 – S6.3) with a high level of significance, indicating that our main results are not sensitive to the method used to merge HR and PD datasets.

### Drivers of the interspecific pattern

In our dataset, body mass varied considerably, ranging from 2.2 kg (*Leopardus tigrinus*) to 163 kg (*Panthera tigris*). Given that larger animals need more resources, they exhibit lower PDs (Damuth, 1981) and require larger home ranges to meet their daily needs (McNab, 1963). Body mass correlated with both HRS and PD (Spearman |*ρ*| > 0.67, Table S9.1), the largest felids having the largest home ranges and the lowest PDs. Even though these results are correlative, it is likely that body mass is a major driver of the HRS– PD relationship between *Felidae* species. However, the statistical effect of PD on HRS remained significant (*p* = 0.042) when accounting for body mass (Table S9.2), suggesting that other factors may drive the interspecific pattern. Determining which mechanisms are at play is beyond the scope of this study, and future research should explore this point deeper.

The relationship between HRS and PD is linear on a log-scale, which was expected since both scale allometrically with body mass. Let us call α the scaling factor between HRS and body mass (i.e. the slope of the linear relationship between HRS and body mass after log-transformation), and δ the scaling factor between PD and body mass. Makarieva, Gorshkov & Li (2005) predicted that α = - δ in a stable ecosystem and α > - δ in a disturbed ecosystem.

This implies that HRS ∝ PD^α/δ^ with a scaling factor α/δ less than or equal to -1. Our interspecific scaling factor (−1.323 ± 0.180) is thus compatible with known allometric relationships between HRS, body mass and PD.

### Drivers of the intraspecific pattern

Within a given species, PD is significantly and negatively associated with HRS, and this effect is not confounded with environmental variables, suggesting that PD may be an inherent driver of HRS. This result is compatible with the conspecific competition hypothesis (H_3_), which predicts that an increase in PD causes an increase in competition, notably competition for space, leading to a reduction in HRS (Sells & Mitchell, 2020). It is also in line with the marked territorial behavior displayed by felids (Bradshaw, 2016). However, whether PD causes a reduction of HRS or HRS causes a reduction of PD, is not fully elucidated, because we could also postulate that HRS regulates PD by inducing the dispersal of the individuals which do not hold a territory (Jones, 1990). One possibility to disentangle these two processes would be to collect longitudinal records on PD and HRS, to see which mechanism comes first in time and how dispersal rates respond to HRS and PD variation.

HRS not only relates to PD, but also to croplands density. We can hypothesise that croplands, along with other human activities to which they are connected (e.g. HPD, HFI, road density, Figure S8.1), induce habitat fragmentation and thus restrict the home range (Hinam & Clair, 2008). Alternatively, anthropised habitats and agricultural lands might provide available resources for felids, such as livestock (Khorozyan, Ghoddousi, Soofi & Waltert, 2015).

However, although croplands density negatively correlated with HRS, it did not show a significant positive correlation with PD in the clustering approach (Figure S8.1a – b), while both correlations are expected under the “habitat quality” hypothesis. In the country approach, only when applying spatio-temporal averaging did some variables associated with HRS (NPP, elevation, and cropland density, Table S6.3) show an opposite relationship with PD (Figure S8.1c – d). Overall, the predictions of the “habitat quality” hypothesis (H_2_) are not robustly verified in our dataset.

Habitat quality is greatly determined by resource density (Balluffi-Fry et al., 2025). However, the variables we used to approximate prey density (NPP, AR, RR) may be too indirect. NPP is a proxy of energy availability for herbivores, which are in turn consumed by felids.

Although we often see a positive correlation between the number of individuals—such as the number of preys items—and both energy availability and species richness, there are also many counterexamples (Storch, Bohdalková & Okie, 2018). Therefore, prey abundance data are needed to directly assess the effect of resource density on HRS and PD. In a preliminary stage, we attempted to take advantage of the TETRAdensity database (Santini et al., 2024) to extract the densities of mammals (Rodentia and Artiodactyla) in the vicinity of our clusters of interest. However, this significantly reduced the number of HRS-PD pairs, as well as the number of species taken into account in the analysis, in such a way that no robust conclusion could be drawn.

It is worth noting that we could not test the body mass hypothesis (H_1_) at the intraspecific level, because we did not have body mass estimates at the population level. This hypothesis is nonetheless plausible, since body mass has been found to scale positively with HRS (Schradin et al., 2010) and negatively with PD (Dunham & Vinyard, 1997) within some Vertebrates species. Further studies are needed to fill this knowledge gap.

Finally, the intraspecific scaling factor between HRS and PD ranged from -0.569 ± 0.201 to - 0.537 ± 0.201, which is weaker in absolute terms than the interspecific factor we found (−1.323 ± 0.180). It is also weaker than -1, which was assumed by Jetz et al. (2004) in the case of exclusive territories. Assuming this difference is significant, it may be interpreted as a “conservative” behaviour exerted by felids, which can retain large home ranges despite increasing PD, theoretically leading to increasing overlaps between territories (Sells & Mitchell, 2020). However, the process of merging two independent datasets can also increase the dispersion of the data points once paired (see the methodological limitations below), so we cannot rule out that our values for the scaling coefficient are underestimated.

### Methodological limitations

One of the main drawbacks of merging different datasets is that merging can increase the variability of the HRS-PD pairs. Indeed, when pairing HRS and PD estimates that are not perfectly aligned in space and time, each HRS estimate may be associated with an underestimated or an overestimated PD. This may result in an additional “noise”, obscuring the true relationship between HRS and PD. Nonetheless, we were still able to capture this pattern in the clustering approach, as well as in the country approach (Tables S6.1 – S6.3), where HRS and PD data points were further apart from each other. This supports the rationale of merging two independent datasets when investigating relationships that can occur at large spatial scales.

Furthermore, we acknowledge that our sampling is unbalanced and that certain species are much more represented than others (Table S3.1). Especially, in the clustering approach, the filtering process was to the detriment of the smallest felids, probably because their dispersal range is smaller than that of large felids. To ensure that the negative relationship between HRS and PD was not due to just a few species, we conducted a sensitivity study using a leave-one-out approach (Table S11.1), removing each species at a time and applying the selected model to these sub-datasets. The results of this sensitivity analysis show that the pattern was not impacted by the removal of any species except for *Panthera onca* in the clustering approach. Note that in the country approach, results were insensitive to the removal of any species (Table S11.1), certainly due to the large number of species initially present in this dataset. Overall, HRS is robustly negatively related to PD in the *Felidae* family at the intraspecific level.

### Perspectives

As an opening statement, our results may have important implications for epidemiology. For example, culling campaigns have been carried out against Eurasian badgers (*Meles meles*) in the UK to reduce the transmission of bovine tuberculosis in the early 2000s. However, this campaign had mixed and contradictory effects, likely because culling induced perturbations of the social structure of badgers, leading to an increase in individual movements and an elevated risk of disease transmission between badgers and cattle (Pope et al., 2007). As infectious diseases can jeopardise endangered species, understanding the drivers of the HRS-PD relationship might be of special importance when dealing with conservation programs (Lafferty & Gerber, 2002). We notably predict that if the “conspecific competition” hypothesis (H_3_) is true, a measure aimed at increasing PD may not increase disease transmission to the same extent, since a mitigating effect would occur along with a reduction of HRS. Still, the precise ecological consequences of our results are yet to be fully understood, leading the path to more ecological studies on the relationship between HRS and PD. We also encourage further similar research on other taxonomic groups, notably non-territorial animals, in order to explore whether a relationship between HRS and PD is universally supported, and whether the underlying mechanisms are conserved across taxa.

## Authors contribution and acknowledgements

Noé Bugaud: merging of HR and PD datasets, statistical analyses, graphical representations, drafting, conceptualisation. Stefano Anile: construction of PD database, draft reviewing, supervision. Arthemis Moraru: construction of HR database, addition of environmental variables, draft reviewing. Sébastien Devillard: supervision, conceptualisation, draft reviewing.

## Data and code availability statement

Home Range, Population Density and auxiliary variables datasets as well as R codes are publicly available in Bugaud, N., Anile, S., Moraru, A., & Devillard, S. (2026). Data and Code for the paper “Home range size and population density are negatively correlated in wild felids globally”. https://doi.org/10.5281/zenodo.20233208

## Data and code availability statement

Home Range, Population Density and auxiliary variables datasets as well as R codes are publicly available at https://doi.org/10.5281/zenodo.20233208.

