## Supplementary materials for "Home range size and population density are negatively correlated in wild felids globally"

### S1 – Simple and spatio-temporal averaging

After grouping HRS and PD records within clusters (resp. countries), we used two methods for pairing HRS and PD estimates within the same cluster. The first was called "simple averaging", and consisted in pairing the average HRS of a cluster (distinguishing by sex and HRS calculation method) with the average PD of the same cluster (Figure S1.1a). The second was called "spatio-temporal averaging", and consisted in pairing each HRS record with a weighted average of all the PD records in the cluster (Figure S1.1b).

We calculated those weights as numbers negatively related to the spatio-temporal distances between HRS and PD data points; they hence simultaneously accounted for:

(1) the spatial distance (*Ds*), which was the real distance (in metres) computed with the *distm* function from the 'geosphere' package (Hijmans, 2010) in R (R Core Team, 2025);

(2) the temporal distance (*Dt*), which was the Euclidean distance between the sampling time of the two data points (HRS and PD):

$$Dt = \sqrt{(PDstudystartyear - HRSstudystartyear)^2 + (PDstudyendyear - HRSstudyendyear)^2}$$

(3) the proportion of overlap between the monitoring periods of both records (*Ov*), so as to assign greater weight when study periods overlapped:

$$Ov = 2 \times \frac{Durationofoverlap}{DurationofPDstudy + DurationofHRSstudy}$$

Eventually, the two distances *Ds* and *Dt* were scaled and the weights were calculated as follows:

$$Weight = 2^{4Ds + Dt + (1 - Ov)}$$

The exponential relationship was implemented to obtain positive values and to emphasise the importance of the closest data points. In addition, the constant 4 was added to give more weight to the spatial distance than the temporal distance, assuming that populations are primarily characterised by their spatial location.

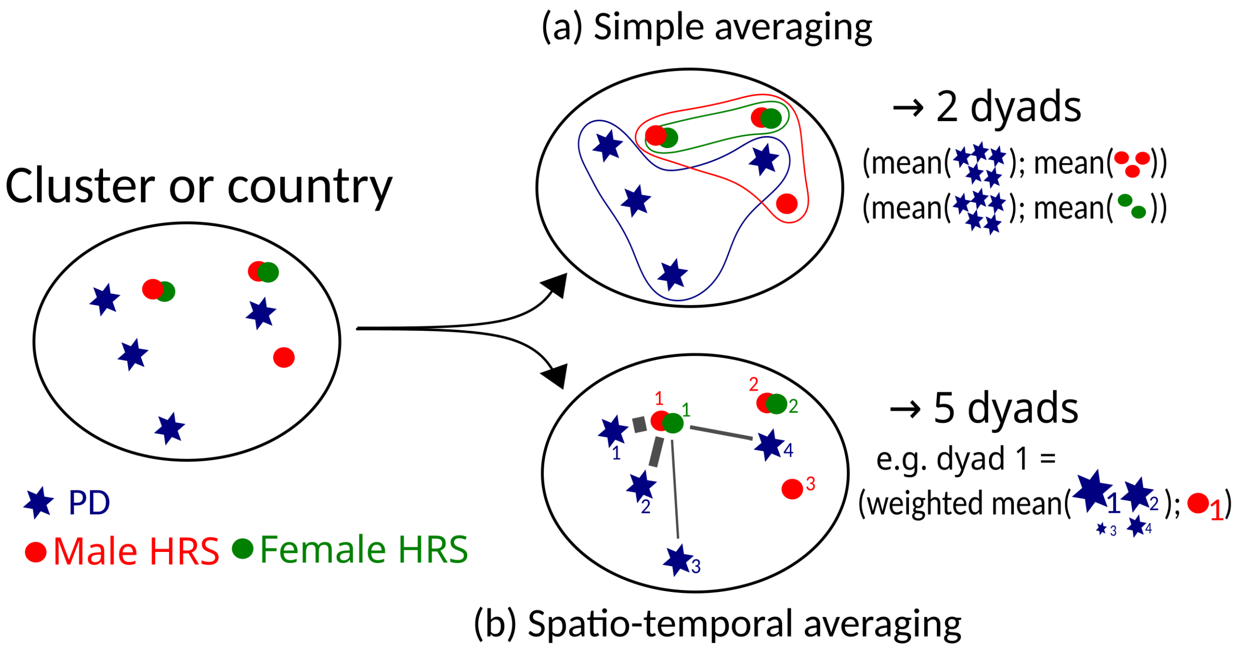

**Figure S1.1.** Illustration of simple averaging **(a)** and spatio-temporal averaging **(b)**. HRS data points represent the mean HRS calculated in a given locality: one for males and one for females. For simplicity, we do not show the distinction between MCP and KDE methods. In (b), the thickness of the grey lines represents the weight, i.e. the contribution of each PD estimate to the final pair.

### S2 – Environmental variables

**Table S2.1.** Environmental variables used in linear mixed-effects models for the intraspecific study.

| Variable name | Source | Comments |
|---|---|---|
| Artiodactyla richness (AR)<br>Rodentia richness (RR) | IUCN 2018<br>Raster maps extracted from https://biodiversitymapping.org/index.php/mammals/ | Spatial resolution: 10x10 km.<br><br>Year: 2018 |
| Elevation* | Hijmans, Barbosa, Ghosh & Mandel, 2024 | Spatial resolution: 1 km (30 arc seconds) aggregated from 90 m resolution data. |
| Net primary productivity (NPP)* | Running & Zhao, 2021 | Spatial resolution: 0.1 degrees (~11x11 km). |
| Human footprint index (HFI)* | Mu et al., 2021 | Spatial resolution: 1 km (30 arc seconds). |
| Human population density (HPD)* | Center For International Earth Science Information Network-CIESIN-Columbia University 2017 | Spatial resolution: 2.5 arc minutes (~4x4 km). |
| Road density* | Meijer et al., 2018 | Spatial resolution: 5 arc minutes (~8x8 km). |
| Croplands density* | Ramankutty, Evan, Monfreda & Foley, 2008 | Spatial resolution: 5 arc minutes (~10 km). |
| Pastures density* | Ramankutty et al., 2008 | Spatial resolution: 5 arc minutes (~10 km). |

* Variables present in the HR dataset used by Moraru, Anile & Devillard, 2026

### S3 – Number of records per species

**Table S3.1.** Number of home range size (HRS) records and population density (PD) records for each species.

| | (i) HR and PD datasets | | | (ii) Clustering approach | | | | | (iii) Country approach | | | | |
|---|---|---|---|---|---|---|---|---|---|---|---|---|---|
| *Species* | PD records | HRS records (individuals) | HRS records (populations) | Clusters | PD records | HRS records (individuals) | HRS/PD pairs (simple averaging) | HRS/PD pairs (spatio-temporal averaging) | Countries | PD records | HRS records (individuals) | HRS/PD pairs (simple averaging) | HRS/PD pairs (spatio-temporal averaging) |
| *Leopardus tigrinus* | 2 | 1 | 1 | | | | | | 1 | 2 | 1 | 1 | 1 |
| *Leopardus guttulus* | 1 | 4 | 4 | | | | | | 1 | 1 | 4 | 4 | 4 |
| *Prionailurus bengalensis* | 18 | 6 | 4 | | | | | | 1 | 2 | 2 | 2 | 2 |
| *Pardofelis marmorata* | 8 | 3 | 2 | | | | | | | | | | |
| *Leopardus wiedii* | 11 | 7 | 3 | | | | | | | | | | |
| *Felis silvestris* | 18 | 11 | 5 | | | | | | | | | | |
| *Otocolobus manul* | 1 | 4 | 2 | 1 | 1 | 4 | 2 | 2 | 1 | 1 | 4 | 2 | 2 |
| *Leopardus geoffroyi* | 3 | 36 | 7 | 1 | 2 | 20 | 4 | 4 | 2 | 3 | 26 | 5 | 5 |
| *Herpailurus yagouaroundi* | 6 | 9 | 9 | | | | | | | | | | |
| *Lynx rufus* | 11 | 156 | 23 | | | | | | 1 | 10 | 156 | 4 | 23 |
| *Lynx canadensis* | 2 | 28 | 9 | | | | | | 2 | 2 | 28 | 3 | 9 |
| *Leptailurus serval* | 21 | 45 | 4 | 1 | 2 | 45 | 4 | 4 | 1 | 13 | 45 | 4 | 4 |
| *Lynx pardinus* | 4 | 10 | 3 | | | | | | 1 | 3 | 10 | 2 | 3 |
| *Leopardus pardalis* | 83 | 42 | 11 | | | | | | 3 | 44 | 34 | 9 | 9 |
| *Neofelis nebulosa* | 15 | 6 | 4 | | | | | | 1 | 2 | 6 | 4 | 4 |
| *Lynx lynx* | 55 | 142 | 34 | 7 | 31 | 91 | 14 | 18 | 5 | 35 | 87 | 11 | 15 |
| *Acinonyx jubatus* | 23 | 15 | 6 | | | | | | 1 | 2 | 10 | 4 | 4 |
| *Panthera pardus* | 312 | 80 | 33 | 12 | 72 | 65 | 27 | 27 | 7 | 231 | 59 | 19 | 23 |
| *Puma concolor* | 41 | 123 | 25 | 1 | 1 | 44 | 2 | 2 | 3 | 13 | 103 | 9 | 17 |
| *Panthera uncia* | 42 | 30 | 12 | 1 | 2 | 22 | 4 | 4 | 1 | 4 | 30 | 4 | 12 |
| *Panthera onca* | 168 | 83 | 55 | 10 | 78 | 60 | 20 | 39 | 5 | 90 | 83 | 12 | 55 |
| *Panthera leo* | 19 | 36 | 7 | 1 | 1 | 21 | 2 | 2 | 2 | 2 | 26 | 4 | 4 |
| *Panthera tigris* | 197 | 145 | 29 | 6 | 78 | 48 | 16 | 18 | 4 | 175 | 145 | 10 | 29 |
| **Total** | **1061** | **1022** | **292** | **41** | **268** | **420** | **95** | **120** | **25** | **635** | **859** | **113** | **225** |

(i) Number of records in the initial PD and HR datasets. The HRS records refer either to individual records, or to population estimates after averaging by location, sex, and the method used to calculate HRS (MCP or KDE).
(ii) Number of observations available for each species after merging with the clustering approach.
(iii) Number of observations available for each species after merging with the country approach. The "country" column corresponds to the number of countries which contain at least one HRS and one PD value for a given species and the "total" refers to the total number of countries represented in the dataset.

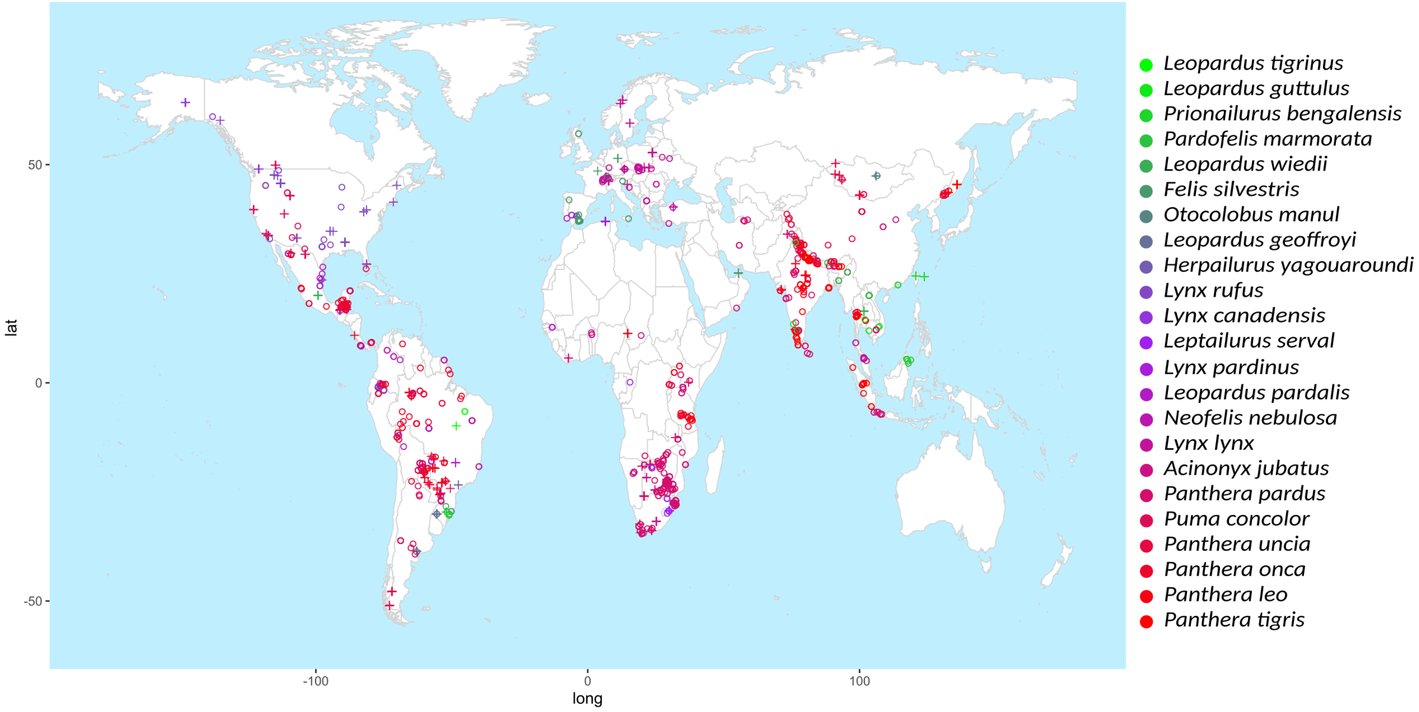

**Figure S3.1.** Geographic positions of HRS (crosses, n = 292) and PD (circles, n = 1061) records for species present in both HR and PD datasets, seen in Mercator projection. Species are listed in order of increasing body mass. The map was generated with the 'ggmap' package (Kahle & Wickham, 2013).

#### S4 – Interspecific relationship with *Lynx pardinus*

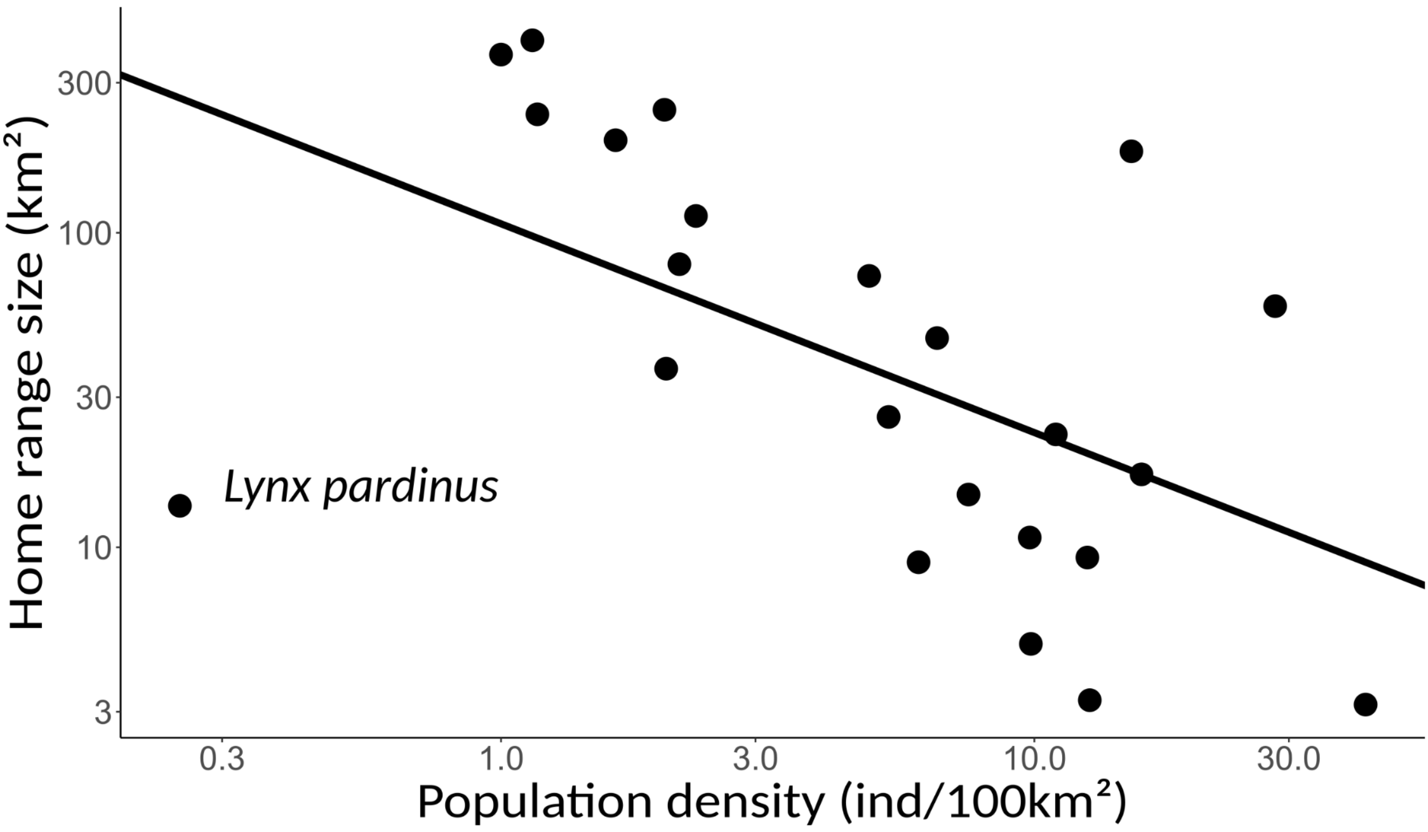

**Figure S4.1.** Interspecific relationship between HRS and PD in Felids, including *Lynx pardinus* (bottom-left data point). Each species was represented by its median HRS and its median PD value across all estimates recorded in HR and PD datasets. We fitted a simple linear regression and estimated a slope of -0.664 with a standard error SE = 0.224 and *p*-value = 0.00747. Residuals were normally distributed (Shapiro test, $p = 0.975$) and had a homogeneous variance (Breush-Pagan test, $p = 0.208$).

### S5 – Clustering approach: selection tables

***Cluster centroid selection***

**Table S5.1.** Number of HRS/PD pairs, species and clusters obtained according to the nature of the data points used as initial centroids in the K-means clustering method.

| | Number of pairs | | Number of species | Number of clusters |
|---|---|---|---|---|
| | Simple averaging | Spatio-temporal averaging | | |
| Clusters based on HRS centroids | 67 | 86 | 9 | 30 |
| Clusters based on PD centroids | 95 | 120 | 10 | 41 |

***Random structure selection***

**Table S5.2.** Clustering approach: random structure selection. The initial full model was: $\log_{10}$(HRS) ~ $\log_{10}$(PD) + Sex + HRS calculation method + NPP + Elevation + AR + RR + Pastures + Croplands + Sex:$\log_{10}$(PD) + Species as a random effect (intercept and/or slope). The selected structure is shown in bold.

| Random effect | Simple averaging | | Spatio-temporal averaging | |
|---|---|---|---|---|
| | df | AICc | df | AICc |
| Slope | 13 | 112.6782 | 13 | 145.2915 |
| Intercept | 13 | 101.8066 | 13 | 143.4113 |
| **Slope + intercept** | **15** | **100.2118** | **15** | **135.0803** |

*df: degree of freedom; AICc, corrected Akaike information criterion.*

***Fixed structure selection***

**Table S5.3.** Clustering approach: fixed structure selection. The five 'best' models and the null model are presented. The initial full model was: $\log_{10}$(HRS) ~ $\log_{10}$(PD) + Sex + HRS calculation method + NPP + Elevation + AR + RR + Pastures + Croplands + Sex:$\log_{10}$(PD), with Species as a random intercept and a random slope. The models retained are shown in bold. N = 10 species.

| Approach | Model | df | logLik | AICc | ΔAICc | Weight |
|---|---|---|---|---|---|---|
| Simple averaging | **HRS ~ Sex + Croplands + PD + HRS calculation method** | **9** | **-23.18** | **66.47** | **0** | **0.4** |
| | **HRS ~ Sex + Croplands + HRS calculation method** | **8** | **-25.27** | **68.21** | **1.74** | **0.17** |
| | HRS ~ Sex + Croplands + PD | 8 | -26.22 | 70.12 | 3.65 | 0.06 |
| | HRS ~ Sex + Croplands + PD + HRS calculation method + Sex:PD | 10 | -23.81 | 70.24 | 3.77 | 0.06 |
| | HRS ~ Sex + Croplands + PD + HRS calculation method + AR | 10 | -24.34 | 71.3 | 4.83 | 0.04 |
| | Null | 5 | -49.83 | 110.34 | 43.87 | 0 |
| Spatio-temporal averaging | **HRS ~ Sex + Croplands + PD + HRS calculation method** | **9** | **-42.82** | **105.27** | **0** | **0.25** |
| | **HRS ~ Sex + Croplands + PD** | **8** | **-44.6** | **106.5** | **1.23** | **0.13** |
| | HRS ~ Sex + Croplands + PD + HRS calculation method + AR | 10 | -42.84 | 107.7 | 2.43 | 0.07 |
| | HRS ~ Sex + Croplands + HRS calculation method | 8 | -45.29 | 107.87 | 2.6 | 0.07 |
| | HRS ~ Sex + Croplands + PD + AR | 9 | -44.34 | 108.32 | 3.05 | 0.05 |
| | Null | 5 | -65.95 | 142.42 | 37.15 | 0 |

*HRS: home range size; PD: population density; NPP: net primary productivity; AR: Artiodactyla richness; RR: Rodentia richness; df: degree of freedom; logLik: log-likelihood; AICc, corrected Akaike information criterion; Weight: weights used for model averaging.*

### S6 – Country approach: model selection and results

***Random structure selection***

**Table S6.1.** Country approach: random structure selection. The initial full model was: $log_{10}$(HRS) ~ $log_{10}$(PD) + Sex + HRS calculation method + NPP + Elevation + AR + RR + Pastures + Croplands + Sex:$log_{10}$(PD) + Species as a random effect (intercept and/or slope). The selected structure is shown in bold.

| Random effect | Simple averaging | | Spatio-temporal averaging | |
|---|---|---|---|---|
| | df | AICc | df | AICc |
| Slope | 13 | 172.9892 | 13 | 297.0452 |
| **Intercept** | **13** | **168.6788** | **13** | **275.9572** |
| Slope + intercept | 15 | 173.8638 | 15 | 279.3453 |

*df: degree of freedom; AICc, corrected Akaike information criterion.*

***Fixed structure selection***

**Table S6.2.** Country approach: fixed structure selection. The five 'best' models and the null model are presented. The initial full model was: $log_{10}$(HRS) ~ $log_{10}$(PD) + Sex + HRS calculation method + NPP + Elevation + AR + RR + Pastures + Croplands + Sex:$log_{10}$(PD), with Species as a random intercept. The models retained are shown in bold. N = 19 species.

| Approach | Model | df | logLik | AICc | ΔAICc | Weight |
|---|---|---|---|---|---|---|
| Simple averaging | **HRS ~ PD + Sex + Croplands + NPP** | **7** | **-60.92** | **136.92** | **0** | **0.59** |
| | HRS ~ PD + Sex + Croplands + NPP + AR | 8 | -61.55 | 140.48 | 3.56 | 0.1 |
| | HRS ~ PD + Sex + Croplands + NPP + Sex:PD | 8 | -61.98 | 141.34 | 4.43 | 0.06 |
| | HRS ~ PD + Sex + Croplands + NPP + Elevation | 8 | -62.47 | 142.33 | 5.41 | 0.04 |
| | HRS ~ PD + Sex + Croplands + NPP + Pastures | 8 | -62.69 | 142.77 | 5.85 | 0.03 |
| | Null | 3 | -96.28 | 198.78 | 61.86 | 0 |
| Spatio-temporal averaging | **HRS ~ PD + Sex + Croplands + NPP + AR + Elevation** | **9** | **-117.17** | **253.18** | **0** | **0.55** |
| | HRS ~ PD + Sex + Croplands + NPP + AR | 8 | -119.59 | 255.84 | 2.66 | 0.15 |
| | HRS ~ PD + Sex + Croplands + NPP + AR + Elevation + Pastures | 10 | -118.45 | 257.92 | 4.74 | 0.05 |
| | HRS ~ PD + Sex + Croplands + NPP + AR + Elevation + Sex:PD | 10 | -118.55 | 258.13 | 4.95 | 0.05 |
| | HRS ~ PD + Sex + Croplands + NPP | 7 | -121.84 | 258.19 | 5.01 | 0.05 |
| | Null | 3 | -178.19 | 362.48 | 109.29 | 0 |

*HRS: home range size; PD: population density; NPP: net primary productivity; AR: Artiodactyla richness; RR: Rodentia richness; df: degree of freedom; logLik: log-likelihood; AICc, corrected Akaike information criterion; Weight: weights used for model averaging.*

***Results***

**Table S6.3.** Country approach: predictors of home range size (HRS). The parameters are those extracted from the best model, with their estimate, standard error and significance. "simple averaging" consists in joining one average HRS and one average population density (PD) per country (n = 113 pairs). "Spatio-temporal averaging" consists in joining each HRS population record with a weighted average of all PD records in the country, weights being related to the spatial and the temporal proximity between both records (n = 225 pairs). N = 19 species.

| | Simple averaging | | | Spatio-temporal averaging | | |
|---|---|---|---|---|---|---|
| Predictor of $\log_{10}$(HRS) | Coefficient estimate | Standard error | *p*-value | Coefficient estimate | Standard error | *p*-value |
| $\log_{10}$(PD) | -0.49011 | 0.09918 | 4.29e-06 | -0.36568 | 0.07195 | 8.28e-07 |
| Sex (male) | 0.34041 | 0.05476 | 1.63e-08 | 0.35759 | 0.04187 | 3.43e-15 |
| Croplands | -0.16649 | 0.04082 | 8.70e-05 | -0.12366 | 0.03353 | 0.000286 |
| NPP | -0.17566 | 0.04524 | 0.00018 | -0.12553 | 0.02541 | 1.60e-06 |
| AR | | | | -0.18132 | 0.04653 | 0.000130 |
| Elevation | | | | 0.12009 | 0.03862 | 0.002127 |
| Species | Random intercept | | | Random intercept | | |

*NPP: net primary productivity; AR: Artiodactyla richness.*

### S7 – Temporal gap between HRS and PD records

$$Mean absolute distance = \frac{mean(|yearStartPD - yearStartHR|) + mean(|yearEndPD - yearEndHR|)}{2}$$

$$Median absolute distance = \frac{median(|yearStartPD - yearStartHR|) + median(|yearEndPD - yearEndHR|)}{2}$$

**Table S7.1.** Temporal gap between HRS and PD records.

| | Mean absolute distance (years) | | Median absolute distance (years) | |
|---|---|---|---|---|
| | Simple averaging | Spatio-temporal averaging | Simple averaging | Spatio-temporal averaging |
| Clustering approach | 10.17 | 9.17 | 8.10 | 5.37 |
| Country approach | 10.99 | 12.41 | 11.00 | 11.75 |

### S8 – Correlations between predictors of HRS

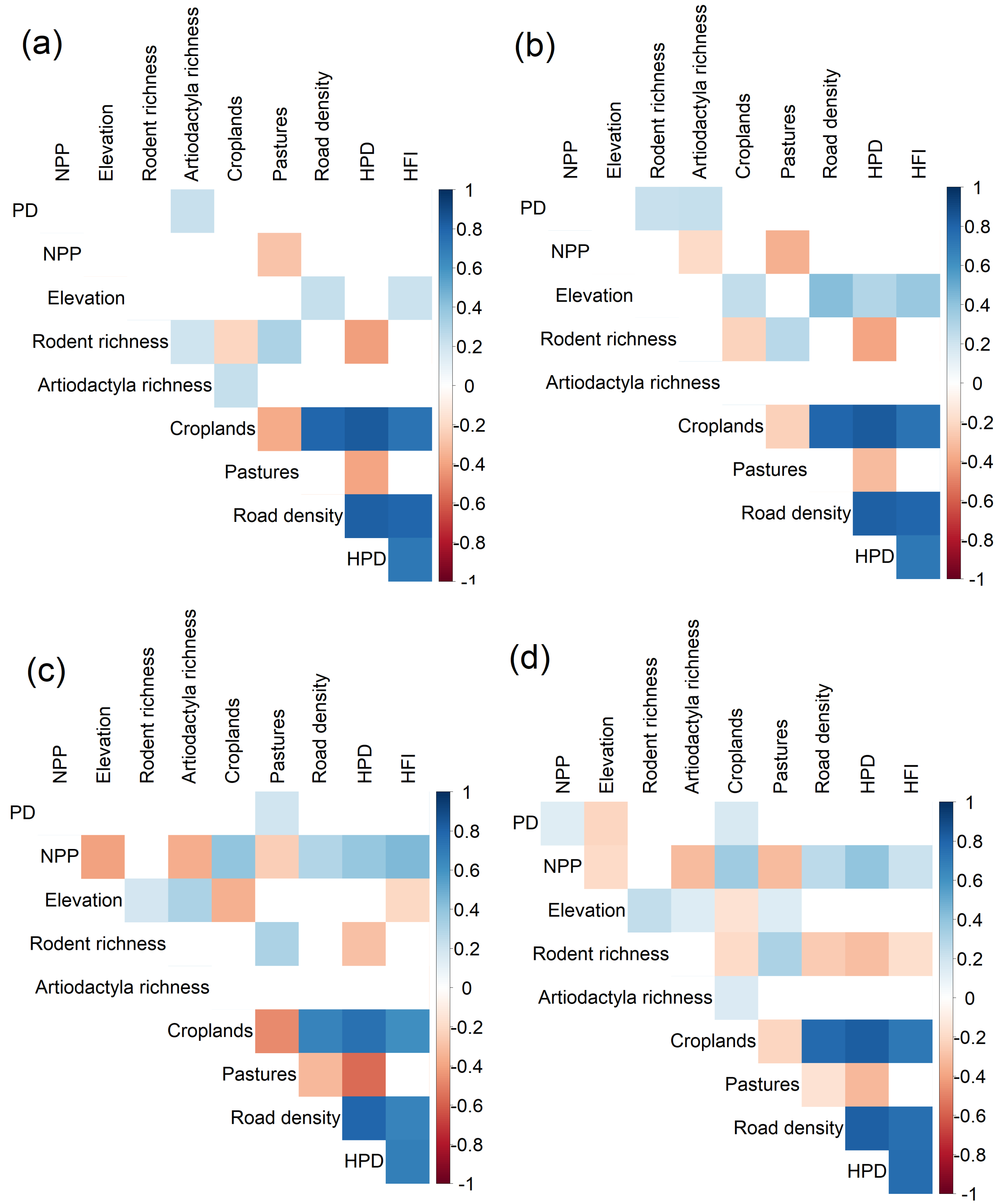

**Figure S8.1.** Spearman correlations $\rho$ between pairs of covariates used to study the interspecific relationship between HRS and PD. Only significant correlations ($p < 0.05$) are shown. **(a)** Clustering approach with simple averaging. **(b)** Clustering approach with spatio-temporal averaging. **(c)** Country approach with simple averaging. **(d)** country approach with spatio-temporal averaging. Variables were considered collinear if $\rho > 0.6$.

### S9 – Interspecific relationship between HRS, PD and body mass

**Table S9.1.** Interspecific spearman correlations $\rho$ between HRS, PD and body mass. Each species was assigned the median HRS and the median PD based on the HR and PD datasets. Each species was also assigned either a specific body mass, or the average mass of males and females (Moraru et al., 2026). *Lynx pardinus* was not included in the analysis.

| | PD | Body mass |
|---|---|---|
| HRS | $\rho$ = -0.722, *p* = 0.0002213 | $\rho$ = 0.767, *p* = 4.668e-05 |
| PD | | $\rho$ = -0.670, *p* = 0.0008727 |

**Table S9.2**. Parameter estimates of the simple linear regression: HRS ~ PD + body mass. There was no significant violation of normality (Shapiro test, *p* = 0.056) or homoscedasticity (Breush-Pagan test, *p* = 0.125) of residuals. The model without interaction between PD and body mass was more parsimonious (ΔAICc = 3.1) than the model with interaction.

| Predictor of $\log_{10}$(HRS) | Estimate | Standard error | *p*-value |
|---|---|---|---|
| $\log_{10}$(PD) | -0.6068 | 0.2781 | 0.04188 |
| $\log_{10}$(body mass) | 0.4829 | 0.2186 | 0.03966 |

### S10 – Model validation plots

#### S10.1 – Interspecific study

To check the assumptions (Zuur, Ieno, Walker, Saveliev & Smith, 2009) of the generalised least square (GLS) model, we:
i) plotted the standardised residuals versus the fitted values and used a Breusch-Pagan test of variance homogeneity, as implemented in the 'performance' package (Lüdecke, Ben-Shachar, Patil, Waggoner & Makowski, 2021);
ii) plotted a histogram of the standardised residuals, and performed a Shapiro test of normality.

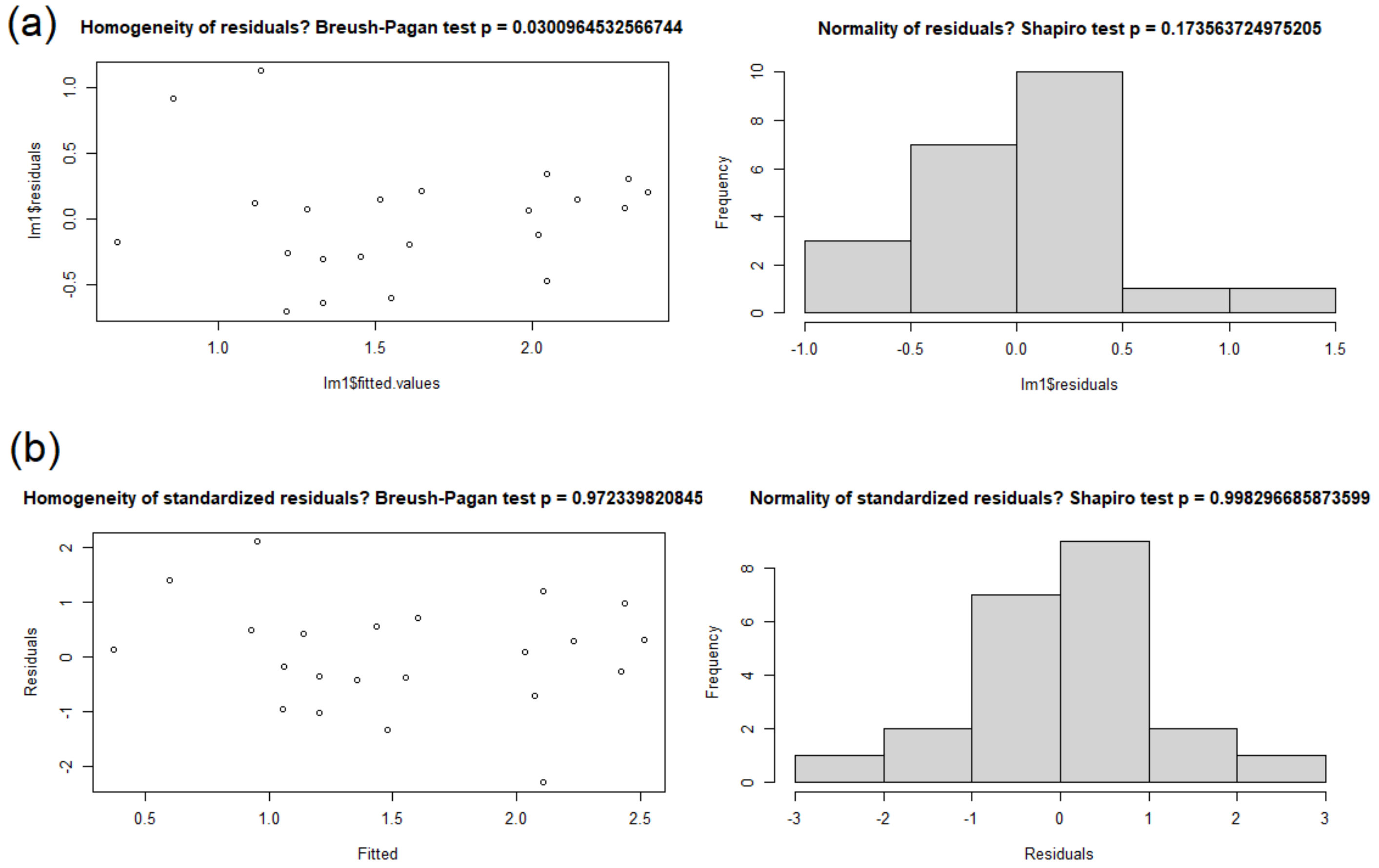

**Figure S10.1.** Validation plots of the simple linear model **(a)** and the generalised least squares (GLS) model **(b)** used for the interspecific analysis. The residuals are non-homogeneous in the simple linear model. All assumptions are respected for the GLS model.

#### S10.2 – Intraspecific study

To check the assumptions (Zuur et al., 2009) of the linear mixed-effect model (LMM), we:
i) plotted the residuals versus the fitted values and used a Breusch-Pagan test of homoscedasticity, as implemented in the 'performance' package (Lüdecke et al., 2021);
ii) plotted a histogram of the residuals, and performed a Shapiro test of normality;
iii) plotted a histogram of the random effects (intercepts and/or slopes), and performed a Shapiro test of normality.

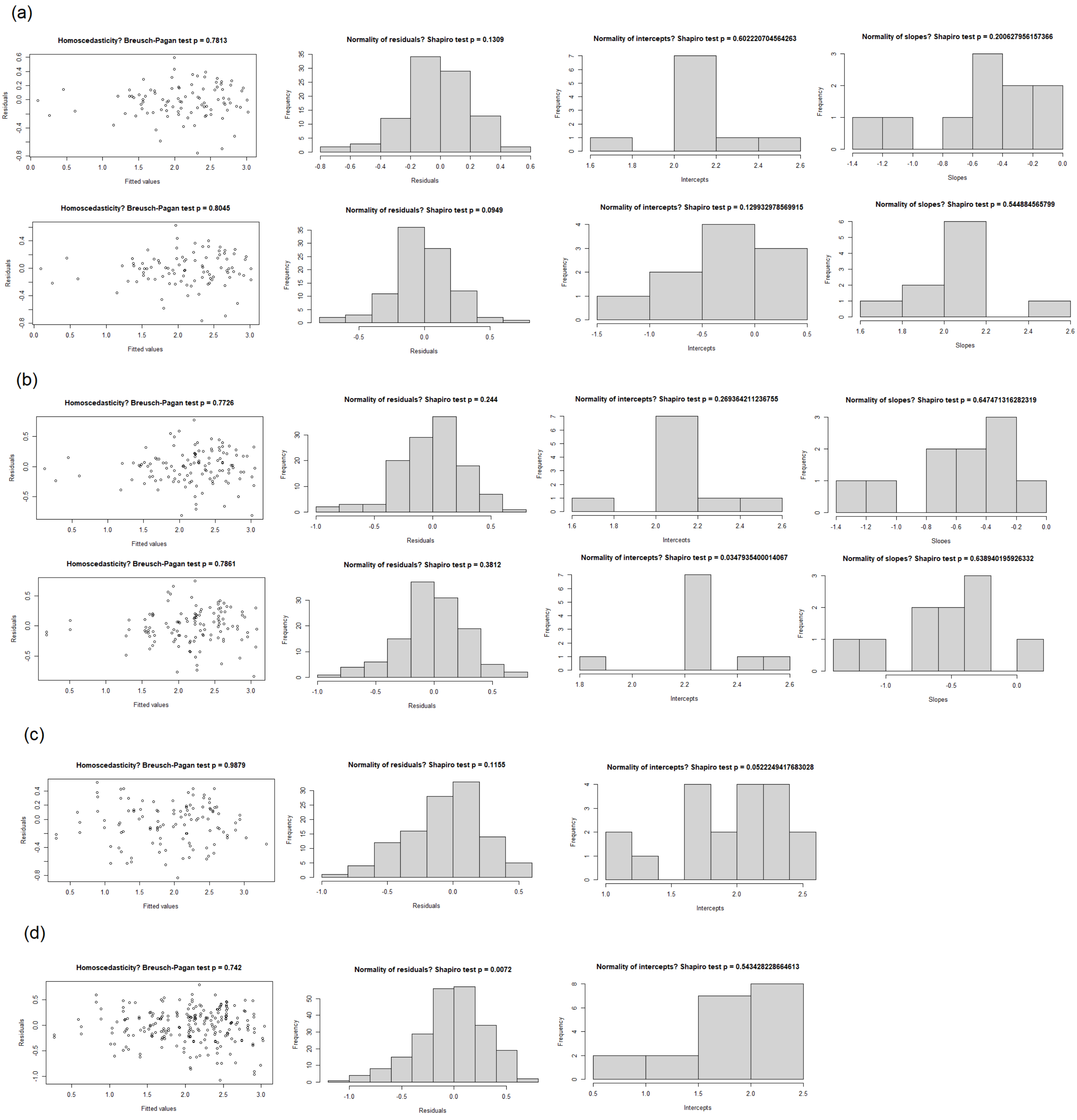

**Figure S10.2.** Validation plots of the mixed-effects models retained, shown in ascending order of their AICc. First column: residuals versus fitted values. Second column: histogram of residuals. Third column: histogram of random intercepts. Fourth column: histogram of random slopes. **(a)** Clustering approach with simple averaging. **(b)** Clustering approach with spatio-temporal averaging. **(c)** Country approach with simple averaging. **(d)** Country approach with spatio-temporal averaging. Overall, the assumptions of the linear mixed-effects models are respected, except the normality of the intercepts in the second 'best' model of the clustering approach with spatio-temporal averaging (b), and the normality of the residuals in the country approach with spatio-temporal averaging (d).

### S11 – Sensitivity analysis

To investigate whether or not the relationship between PD and HRS we found on the full dataset is influenced by the unbalanced representation of species in the dataset, we used a leave-one-out approach as sensitivity analysis. Starting from the best supported model for each dataset, we removed one species at a time and assessed both the value and the significance of the PD slope (Table S11). We summarised the sensitivity analysis by the proportion of models showing a significant relationship between PD and HRS as well as by the coefficient of variation of the slope value over all models (Table S11).

For the clustering approach, 14 out of 20 (70%) models showed a significant relationship between PD and HRS (Table S11). When removing *Panthera onca* in the clustering approach, the relationship between HRS and PD was only marginally significant ($p < 0.105$). However, overall, the relationship remained negative, the *p*-values remained low, and the slope value was largely constant, with coefficients of variation equal to 0.141 and 0.152 for simple and spatio-temporal averaging, respectively. Overall our sensitivity analysis on the clustering approach showed strong support for the relationship between PD and HRS.

For the country approach, the slope between PD and HRS was always significant regardless of the species removed, suggesting the pattern is strongly supported by the data. The absolute coefficient of variation of the slope value was low for simple (0.109) and spatio-temporal averaging (0.126).

**Table S11.1.** Sensitivity analysis: effect of PD on HRS when one species is removed from the dataset. Each cell displays the estimate of the PD effect ± its standard error, and the number of pairs removed (n) in parentheses. * $p < 0.05$; ** $p < 0.01$; *** $p < 0.001$.

| | Clustering approach | | Country approach | |
|---|---|---|---|---|
| Species removed | Simple averaging | Spatio-temporal averaging | Simple averaging | Spatio-temporal averaging |
| *Leopardus tigrinus* | Species not included in this dataset | Species not included in this dataset | -0.484 ± 0.096 (n = 1) *** | -0.363 ± 0.072 (n = 1) *** |
| *Leopardus guttulus* | Species not included in this dataset | Species not included in this dataset | -0.485 ± 0.1 (n = 4) *** | -0.363 ± 0.072 (n = 4) *** |
| *Prionailurus bengalensis* | Species not included in this dataset | Species not included in this dataset | -0.478 ± 0.103 (n = 2) *** | -0.358 ± 0.073 (n = 2) *** |
| *Otocolobus manul* | -0.568 ± 0.218 (n = 2), *p* = 0.053 | -0.61 ± 0.214 (n = 2) * | -0.497 ± 0.1 (n = 2) *** | -0.367 ± 0.073 (n = 2) *** |
| *Leopardus geoffroyi* | -0.401 ± 0.106 (n = 4) * | -0.418 ± 0.152 (n = 4) * | -0.447 ± 0.098 (n = 5) *** | -0.348 ± 0.072 (n = 5) *** |
| *Lynx rufus* | Species not included in this dataset | Species not included in this dataset | -0.495 ± 0.102 (n = 4) *** | -0.348 ± 0.078 (n = 23) *** |
| *Lynx canadensis* | Species not included in this dataset | Species not included in this dataset | -0.528 ± 0.105 (n = 3) *** | -0.384 ± 0.075 (n = 9) *** |
| *Leptailurus serval* | -0.471 ± 0.203 (n = 4), *p* = 0.06 | -0.505 ± 0.203 (n = 4) * | -0.491 ± 0.102 (n = 4) *** | -0.363 ± 0.072 (n = 4) *** |
| *Lynx pardinus* | Species not included in this dataset | Species not included in this dataset | -0.592 ± 0.097 (n = 2) *** | -0.407 ± 0.073 (n = 3) *** |
| *Leopardus pardalis* | Species not included in this dataset | Species not included in this dataset | -0.332 ± 0.101 (n = 9) ** | -0.292 ± 0.071 (n = 9) *** |
| *Neofelis nebulosa* | Species not included in | Species not included in | -0.488 ± 0.101 (n = 4) *** | -0.364 ± 0.073 (n = 4) *** |

| | | | | |
|---|---|---|---|---|
| | this dataset | this dataset | | |
| *Lynx lynx* | -0.596 ± 0.221 (n = 14) * | -0.644 ± 0.217 (n = 18) * | -0.475 ± 0.107 (n = 11) *** | -0.37 ± 0.078 (n = 15) *** |
| *Acinonyx jubatus* | Species not included in this dataset | Species not included in this dataset | -0.489 ± 0.103 (n = 4) *** | -0.358 ± 0.073 (n = 4) *** |
| *Panthera uncia* | -0.517 ± 0.192 (n = 4) * | -0.544 ± 0.194 (n = 4) * | -0.486 ± 0.1 (n = 4) *** | -0.366 ± 0.071 (n = 12) *** |
| *Panthera pardus* | -0.651 ± 0.222 (n = 27) * | -0.679 ± 0.223 (n = 27) * | -0.587 ± 0.093 (n = 19) *** | -0.53 ± 0.073 (n = 23) *** |
| *Puma concolor* | -0.561 ± 0.211 (n = 2), *p* = 0.055 | -0.599 ± 0.214 (n = 2) * | -0.479 ± 0.107 (n = 9) *** | -0.369 ± 0.073 (n = 17) *** |
| *Panthera onca* | -0.457 ± 0.217 (n = 20), *p* = 0.098 | -0.443 ± 0.211 (n = 39), *p* = 0.105 | -0.54 ± 0.105 (n = 12) *** | -0.41 ± 0.085 (n = 55) *** |
| *Panthera leo* | -0.594 ± 0.203 (n = 2) * | -0.609 ± 0.206 (n = 2) * | -0.491 ± 0.101 (n = 4) *** | -0.355 ± 0.072 (n = 4) *** |
| *Panthera tigris* | -0.556 ± 0.232 (n = 16), *p* = 0.058 | -0.604 ± 0.23 (n = 18) * | -0.472 ± 0.113 (n = 10) *** | -0.319 ± 0.085 (n = 29) *** |
| Absolute coefficient of variation | 0.141 | 0.152 | 0.109 | 0.126 |

Note: for each approach, we used the same random and fixed structures as the best model fitted on the dataset containing every species. In the clustering approach, the model was: *HRS ~ Sex + Croplands + PD + HRS calculation method*; species were random intercepts and slopes. In the country approach, the models were: *HRS ~ PD + Sex + Croplands + NPP* (simple averaging), and *HRS ~ PD + Sex + Croplands + NPP + AR + Elevation* (spatio-temporal averaging); species were random intercepts.

Coefficient of variation = estimate standard error / estimate mean.
